# Eltonian niche modelling: applying joint hierarchical niche models to ecological networks

**DOI:** 10.1101/2024.08.14.607929

**Authors:** D. Matthias Dehling, Hao Ran Lai, Daniel B. Stouffer

## Abstract

There is currently a dichotomy in the modelling of Grinnellian and Eltonian niches. Despite similar underlying data, Grinnellian niches are modelled with species-distribution models (SDMs), whereas Eltonian niches are modelled with ecological-network analysis, mainly because the sparsity of species-interaction data prevents the application of SDMs to Eltonian-niche modelling. Here, we propose to adapt recently developed joint species distribution models (JSDMs) to data on ecological networks, functional traits, and phylogenies to model species’ Eltonian niches. JSDMs overcome sparsity and improve predictions for individual species by considering non-independent relationships among co-occurring species; this unique ability makes them particularly suited for sparse datasets such as ecological networks. Our Eltonian JSDMs reveal strong relationships between species’ Eltonian niches and their functional traits and phylogeny. Moreover, we demonstrate that JSDMs can accurately predict the interactions of species for which no empirical interaction data are available, based solely on their functional traits. This facilitates prediction of new interactions in communities with altered composition, e.g. following climate-change induced local extinctions or species introductions. The high interpretability of Eltonian JSDMs will provide unique insights into mechanisms underlying species interactions and the potential impacts of environmental changes and invasive species on species interactions in ecological communities.

## Introduction

There is currently limited methodological overlap between the modelling of Grinnellian and Eltonian niches. *Grinnellian niches*—the range of environmental conditions under which species are known to occur—are commonly modelled with species distribution models (SDMs). SDMs have become a ubiquitous tool in community and macroecology to study the relationship between environmental conditions and species’ geographic ranges and to predict potential future geographic ranges under altered environmental conditions (Elith & Leathwick 2009), with a growing number of methodological approaches (Guisan et al. 2017; Ovaskainen & Abrego 2020). In contrast, *Eltonian niches*—species’ resource use and interactions with other species (Elton 1927)—have classically been studied via ecological networks, such as food webs (Bascompte & Jordano 2014; Dehling 2023), with recent approaches also combining networks with functional traits to describe species’ Eltonian niches using multivariate statistics (Caron et al. 2022; Dehling et al. 2016, 2021, 2022a, b; Junker et al. 2013). This separation (see Björk et al. 2018; Facon et al. 2021; Opedal & Hegland 2020 for exceptions), however, belies the fact that both approaches are based on data of a similar structure: a matrix of species occurrences (site × species for Grinnellian, species × species for Eltonian) combined with environment variables describing the sites where species occur (Grinnellian niche) or characteristics describing a species’ resources or interaction partners (Eltonian niche), respectively (Fig. 1). The similar data structure should make it straightforward to apply modelling approaches geared toward Grinnellian niches to the statistical modelling of Eltonian niches, e.g. to model the current and potential future resource use of a species (Dehling & Stouffer 2018).

**Fig. 1.**
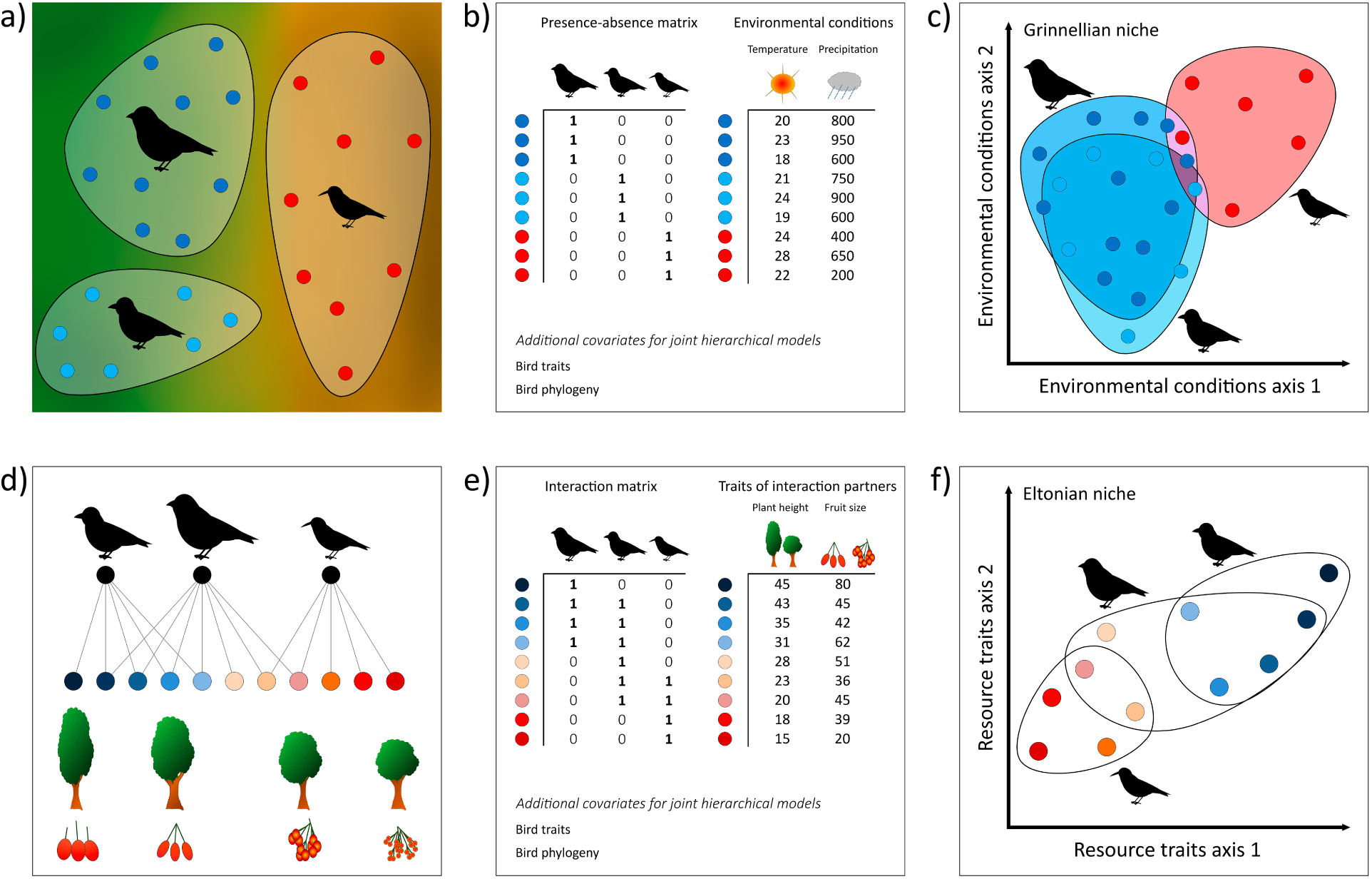
Parallels between the modelling of Grinnellian and Eltonian niches. (a) Grinnellian niches are modelled by assessing the environmental conditions characterizing species’ ranges, either across the entire range or at point samples of known occurrences. (b) Typically, the occurrence of species across the study area is recorded in a presence–absence matrix; the environmental conditions at the different sites are recorded as a site × environment matrix. Grinnellian niches are then modelled based on the environmental conditions found within a species’ range. In joint hierarchical niche models, additional data such as the species’ functional traits or their phylogenetic relationships can be taken into account to improve the modelling of Grinnellian niches. (c) Grinnellian niches can then be plotted in a multidimensional space representing the differences in environmental conditions found at the sites at which species occur. (d) In contrast, Eltonian niches are modelled by assessing species’ resource use or interactions with other species, usually recorded in an interaction network, and the traits of the resources with which each species interacts. (e) Typically, the resource use of different consumers is recorded in a consumer × resource interaction network; the functional traits of resources are recorded in a resource × trait matrix. These two matrices are analogous to the presence–absence matrix and site × environment matrix in (b), respectively. Eltonian niches are then modelled based on the traits of the resources with which a consumer interacts. In joint hierarchical niche models, additional data such as the consumers’ functional traits or their phylogenetic relationships can be taken into account to improve the modelling of Eltonian niches. (f) Eltonian niches can then be plotted in a multidimensional space representing the differences in the traits of the resources used.

The major obstacle that has so far precluded the application of SDMs to the modelling of Eltonian niches lies in the availability of data. While data for modelling Grinnellian niches (i.e., species occurrences and environmental conditions) are often available from large-scale datasets, data for modelling Eltonian niches have remained scarce due to the difficulty of sampling both species interactions and functional traits; this is known as the “Eltonian shortfall” (Peterson et al. 2011). As a result, data on species interactions are much less complete, especially for species that are rare or difficult to observe, which has made it very challenging to apply frameworks based on single-species SDMs to the modelling of species’ Eltonian niches.

Recently introduced joint hierarchical niche models (most commonly formulated as joint species distribution models, JSDMs) aim to improve niche modelling by simultaneously quantifying the niches of all species in a single model and considering non-independent relationships among co-occurring species, such as trait–niche relationships or species’ phylogenetic relatedness (Ovaskainen & Soininen 2011). By “borrowing” data from ecologically or phylogenetically similar species, these models greatly improve predictions for individual species, especially for those species for which limited empirical data are available (Norberg et al. 2019; Ovaskainen et al. 2017). This unique ability of joint hierarchical niche models makes them particularly suited for datasets that include species with incomplete sampling, which is usually the case for ecological networks (Jordano 2016). We therefore propose here the adaptation of the JSDM framework to data from ecological networks, functional traits, and phylogenies to model species’ Eltonian niches.

Conceptually, our approach builds on the framework of describing a species’ Eltonian niche via the traits of its interaction partners (Dehling & Stouffer 2018; Dehling et al. 2022b; Elton 1927). Depending on the ecological process under study, these traits might describe, for example, the prey of a predator, the hosts of a parasite, the flowers visited by a pollinator, or the fruit consumed by a seed disperser. For simplicity, we will hereafter describe the Eltonian niche in all types of interactions as the traits of the *resources* used by a *consumer*. Methodologically, the Eltonian niche modelling approach is based on the joint species distribution modelling framework primarily developed for the Grinnellian niche (Niku et al. 2021; Ovaskainen et al. 2017; Warton et al. 2015), which usually takes the form of a multivariate hierarchical generalized linear latent-variable mixed-effects model. The modelling of Eltonian niches requires data that are similar to those for the modelling of Grinnellian niches, but it involves the following changes (Fig. 1): instead of (i) a matrix describing species occurrences across sites (site × species) and (ii) a matrix describing the environmental conditions across sites (site × environment), Eltonian niche modelling uses (i) the matrix representation of an interaction network describing the consumers’ interactions across resources (resource × consumer) and (ii) a matrix describing the traits across resources (resource × trait). The interaction matrix can be represented either qualitatively (presence–absence) or quantitatively (number of interactions); we focus on the former in this study. Hence, the model assesses the Eltonian niche of consumers based on the traits of the resources that they use.

Eltonian niche modelling can be applied to different types of interactions (predator–prey, plant–pollinator, host–parasite, etc.). Here, we demonstrate our Eltonian niche model on a community of frugivorous bird species (consumers) interacting with fleshy-fruited tree species (resources) using interaction and trait data from a plant–bird seed-dispersal network in the tropical Andes. This allows simultaneous niche modelling of multiple bird species while taking into account co-variation in resource use between them. Moreover, to improve the joint modelling of the birds’ Eltonian niches, we include both functional traits and a phylogeny of the bird species, building on previous work that has demonstrated the strong role of functional traits in determining both realized species interactions and “forbidden links” (physically or ecologically impossible interactions; Olesen et al. 2010, Dehling et al. 2014b, Vizentin-Bugoni et al. 2014). Consequently, inferred interaction coefficients between consumer traits and resource traits can be used to test specific hypotheses about trait matching between consumers and resources. Finally, we demonstrate that joint hierarchical niche models can even be used to predict interactions between consumers and resources for which there are no empirical interaction data available—solely based on the matching of their functional traits—making it straightforward to model new species interactions in assemblages with altered species composition.

## Material and Methods

### Networks

We used an interaction network between frugivorous birds and fleshy-fruited plants sampled in a montane rainforest in the Andes of south-east Peru (Manu Biosphere Reserve; 13.1°S, 71.6°W, 1500 m elevation; Dehling et al. 2014b). The network was sampled over a year with a total sampling effort of 960 h, and consisted of 4,988 interaction events between 61 bird and 52 plant species (further details on network sampling and robustness in Dehling et al. 2014b). The network includes observed interactions between 398 pairs of birds and plants (marked as “1” in the matrix). Rather than treat all 2,774 unobserved interactions as true absences, we used bird and plant trait data (described below) to differentiate between unobserved and unlikely interactions (marked as “0”, 404 species pairs) vs unobserved but possible interactions (marked as “NA”, 2,370 species pairs). See Box 1 for more details.

### Plant traits

We included plant functional traits as predictors to assess how much of the variance in bird–plant interactions is determined by resource traits (i.e., by the birds’ Eltonian niches). The functional traits for all plant species in the networks were collected in the field at the same time as the network observations and included three morphological traits previously shown to influence the interaction between fleshy-fruited plants and frugivorous birds (Bender et al. 2018; Dehling et al. 2014b): fruit diameter (cm), plant height (m), and crop mass (g; the total amount of fruit offered by a tree, calculated as mean number of fruits per plant × mean fruit mass). We used species-level average trait values in the analyses. We also included a quadratic term for fruit diameter because bird species’ niche optima are likely to be at intermediate values.

### Bird traits and bird phylogeny

We used three functional traits shown previously to influence the foraging behaviour of birds because of their strong match with plant traits (Bender et al. 2018; Dehling et al. 2014b): bill width (cm), Kipp’s index (dimensionless; Dehling et al. 2014a), and body mass (g; Dunning 2007, Tobias et al. 2022). Kipp’s index is a measure for the roundedness of the wing and measured on the folded wing as the ratio between the distance from the tip of the first secondary to the wing tip, and the full length of the folded wing. We used species-level average trait values in the analyses. All traits except body mass were measured across several specimens collected close to the network site to assure that traits correspond to the observed interactions (Dehling et al. 2014a); average body mass was taken from the literature (Dunning 2007). We built a phylogeny of all birds in the network as a consensus tree of 1,000 dated phylogenetic trees from birdtree.org (Jetz et al. 2012, 2014) using the phylogenetic backbone by Hackett et al. (2008).

### Model fitting

Leveraging the aforementioned empirical data, we fit one model without including any bird co-variates (model 1) and three additional models with different combinations of bird co-variates: only bird functional traits (model 2), only bird phylogeny (model 3), and both bird functional traits and bird phylogeny (model 4; Table 1). For our analyses, we used the package Hmsc v3.0 (Tikhonov et al. 2020b) in R 4.21 (R Core Team 2022). For all models, we used the default priors in Hmsc (Ovaskainen et al. 2017). For each model, we ran four chains of Markov Chain Monte Carlo (MCMC) with 1,000,000 samples and a burn-in of 200,000 steps. We applied thinning (1,000 step intervals) and ended up with 1,000 posterior samples per chain (4,000 samples per model). We examined model convergence with effective sample sizes and potential scale reduction factors. To compare the different models, we calculated WAIC (Watanabe 2013), and we evaluated in-sample explanatory power by AUC and coefficient of discrimination (Tjur’s *R*^2^). We used two-fold cross-validation and calculated AUC and Tjur’s *R*^2^ to evaluate the out-of-sample predictive power of the models. We also used the computeVariancePartitioning function from the Hmsc package to assess the variance in bird–plant interactions determined by plant traits (associated with the regression slope *β* in models 1–4), the variance in the birds’ Eltonian niches determined by bird traits (associated with *γ* in models 2 & 4), and the phylogenetic signal in Eltonian niches (associated with *ρ* in models 3 & 4) (Table 1, Appendix 2). In order to assess the influence of bird and plant traits on interaction probabilities, we plotted the model-predicted interaction probabilities between birds and plants along gradients of the each plant trait for representative values of each bird trait (Fig. 2).

**Fig. 2.**
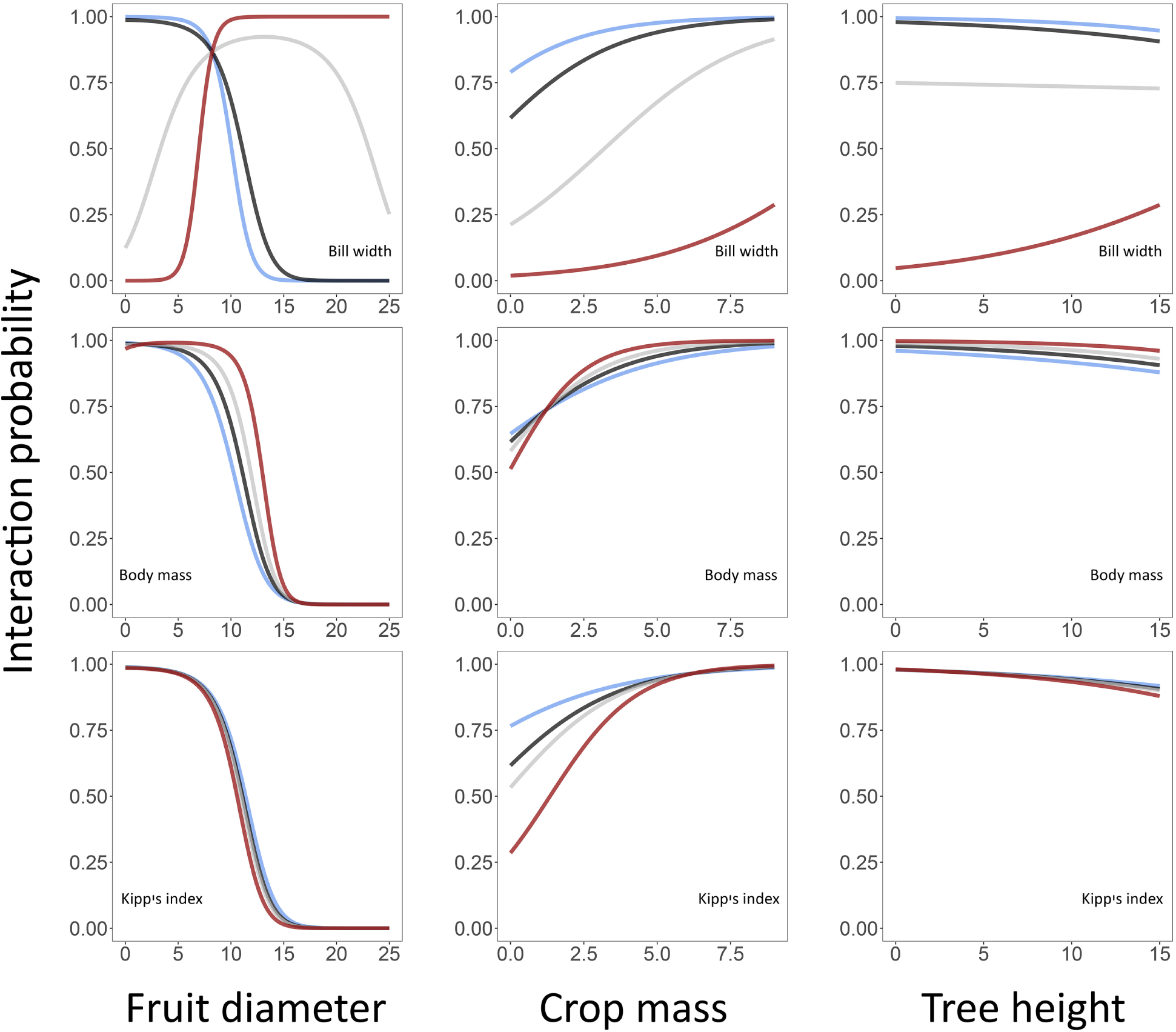
Interaction probabilities between frugivorous birds and fleshy-fruited plants determined by bird and plant traits. Bird traits (rows) include bill width, body mass, and Kipp’s index; plant traits (columns) include fruit diameter, crop mass, and tree height. Posterior-mean interaction probabilities predicted by model 2 (*y*-axis) are shown along a gradient of a focal plant trait (*x*-axis) for four hypothetical bird species with different values of a focal bird trait (blue, 0.025 percentile; black, 0.5 percentile; grey, midpoint between the 0.025 and the 0.975 percentiles; red, 0.975 percentile). For all predictions, non-focal plant traits and non-focal bird traits were set to the respective community median.

**Table 1.** Explanatory and predictive power, explained variance, and phylogenetic signal in Eltonian niches of four joint hierarchical models describing the interactions between 61 frugivorous birds and 52 fleshy-fruited plants. Eltonian niches of bird species are described by the functional traits of the plants with which the birds interact. The joint hierarchical models relate the birds’ Eltonian niches to different combinations of co-variates (no co-variates, bird traits, bird phylogeny, bird traits + bird phylogeny). Percent variance explained describes the mean variance in bird–plant interactions explained by plant traits and the mean variance in Eltonian niches explained by bird traits. The phylogenetic signal in the birds Eltonian niches is given as the 95% interval and median value.

| Model | Co-variates | WAIC | Explanatory power |  | Predictive power |  | Percent variance explained |  | Phylogenetic signal<br>bird traits |
| --- | --- | --- | --- | --- | --- | --- | --- | --- | --- |
|  |  |  | AUC | Tjur R<sup>2</sup> | AUC | Tjur R<sup>2</sup> | bird-plant interactions<br>by plant traits | Eltonian niches<br>by bird traits |  |
| 1 | no co-variates | 2.08 | 1 | 0.87 | 0.92 | 0.65 | 99.6 | - | - |
| 2 | bird traits | 2.24 | 1 | 0.86 | 0.95 | 0.70 | 99.5 | 90.3 | - |
| 3 | bird phylogeny | 1.69 | 1 | 0.90 | 0.95 | 0.66 | 99.8 | - | 0.76–1 (0.97) |
| 4 | bird traits + bird phylogeny | 1.95 | 1 | 0.90 | 0.94 | 0.69 | 99.6 | 70.2 | 0.70–1 (0.95) |

### Predicting new or unobserved species interactions

To test the ability of our approach to model new or unobserved interactions based on proxies such as species’ functional traits, we removed all observed interactions in the empirical network for three species that represent different bird lineages with different morphology (Blue-necked Tanager *Tangara cyanicollis*, Passeriformes: Thraupidae, a common species; Andean Guan *Penelope montagnii*, Galliformes: Cracidae, an uncommon species; Crested Quetzal *Pharomachrus antisianus*, Trogoniformes: Trogonidae, a rare species). For these species, we changed all “1” to “NA” in the interaction matrix (see Box 1). Our reduced interaction network thus contained observed interactions between 58 bird species and the original 52 plant species. We then repeated model 2 (including bird functional traits) by re-fitting it on the data set consisting of the reduced interaction network, but complete consumer × trait (bird traits) and resource × trait (plant traits) data set. We chose model 2 to illustrate the modelling of species interactions from functional traits, as it provides a more intuitive interpretation than modelling interactions from phylogenies. The Eltonian niches for the three removed bird species were thus predicted based only on their functional traits and the inferred Eltonian niches of the remaining bird species. We used correlations to compare the Eltonian niches of the three bird species modelled based on their observed interactions with those predicted based only on their functional traits. For completeness, we repeated this procedure for each bird species in the network in turn, that is, only removing the observed interactions for one bird species at a time (Appendix 1, Fig. S3).

## Results

### Modelling bird–plant interactions via Eltonian niches of birds

The explanatory power of all models (i.e., the explained variance in bird–plant interactions) was very high (AUC: 1, Tjur’s *R*^2^: 0.86–0.90; Table 1). The predictive power of all models, based on two-fold cross validation, was also high, but slightly lower than the explanatory power (AUC 0.65–0.70, Tjur’s *R*^2^: 0.65–0.70, Table 1). Plant traits explained almost all the variance in the bird–plant interactions (Table 1). In turn, bird traits explained 90.3 (model 2) and 70.2 (model 4) percent of the variance in the birds’ Eltonian niches, and there was also a strong phylogenetic signal in the Eltonian niches (model 3: 95% credible interval 0.76–1, median 0.97; model 4: 0.70–1, median 0.95); that is, birds with similar morphology and birds that were more closely related to each other tended to have more similar Eltonian niches.

Unobserved empirical interactions that we determined as possible (“NA”) tended to show a high predicted probability in the models, whereas those determined as unlikely were corroborated as unlikely in the models (Appendix 1, Fig. S1). The resulting high interaction probabilities for most of these unobserved interactions that we identified as possible suggest that treating those interactions as absences would have led to a misrepresentation of the species’ Eltonian niches.

### Trait matching between birds and plants

Models 2 and 4 revealed similar sets of matching bird and plant traits that influenced the interaction between birds and plants, including a humpshaped relationship between bill width and fruit diameter, and between body mass and fruit diameter (Fig. 2, Appendix 1, Fig. S2).

### Predicting species interactions from functional traits

The Eltonian niches of the three bird species (*Tangara cyanicollis*, *Penelope montagnii* and *Pharomachrus antisianus*) predicted based only on their functional traits were very similar to the Eltonian niches modelled based on their empirical interactions in model 2 (correlation of interaction probabilities, *Tangara cyanicollis*, *R*^2^ = 0.99, *p <* 0.001; *Penelope montagnii*, *R*^2^ = 0.94, *p <* 0.001; *Pharomachrus antisianus*, *R*^2^ = 0.98, *p <* 0.001; Fig. 3). The correlations were generally even stronger when we predicted interactions for one bird species at a time (*R*^2^ 0.72–1.0, median 0.995; Appendix 1, Fig. S3).

**Fig. 3.**
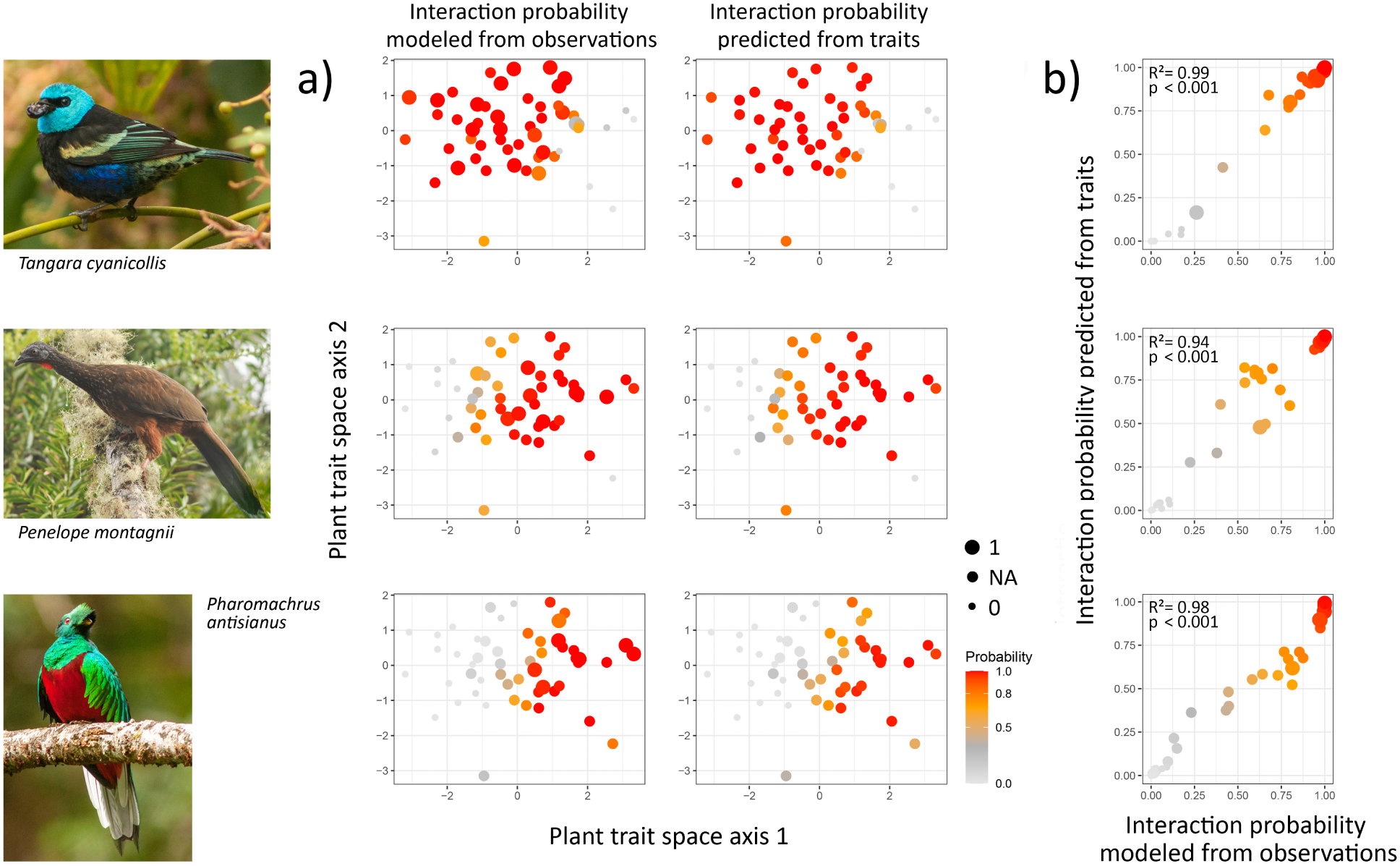
Using joint hierarchical niche models to predict unobserved interactions based on proxies. (a) The Eltonian niches of three bird species in a multi-dimensional plant trait space (only first two axes shown). Plant species are projected into a multidimensional trait space using PCoA. The color of the dots corresponds to the predicted interaction probability between a bird species and a plant species in the network using model 2. For each bird species, interaction probabilities were modelled once from observed interactions (left) and, after removing all observed interactions from the network, predicted based solely on their functional traits and the relationship between traits and Eltonian niches of the other bird species in the network (right). Interactions in the network were marked as observed (1), unobserved but likely (NA), and unobserved and unlikely (0; see Box 1 for details), differences are represented by dot size. (b) Correlations between the interaction probabilities modelled from observed interactions and those predicted based on birds’ functional traits for the three bird species. Bird images by DMD.

## Discussion

Our example using bird–plant interactions demonstrates that joint hierarchical niche models can be used to infer species’ Eltonian niches from ecological network data. The joint models revealed strong influences of plant traits, bird traits, and bird phylogeny on the birds’ Eltonian niches. They also identified generalized interaction trends across a large number of consumer and resource species using a small number of matching traits. Furthermore, the unique ability of joint models to use data from one set of species when making predictions about the niches of new, out-of-sample species demonstrates an elegant way to model previously unobserved interactions in the network.

Species interactions between frugivorous birds and fleshy-fruited plants were clearly captured by the birds’ Eltonian niches (i.e., the range of trait combinations of the plant species with which each bird interacts), which in turn were influenced by the morphology of the bird species. The strong interplay between bird and plant traits on the probability of bird–plant interactions corroborates the role of trait matching—the relationship between a consumer’s traits and the traits of its resources—on species interactions (Caron et al. 2022; Dehling et al. 2014b, 2016; Eklöf et al. 2013; Peralta et al. 2020a, b; Sonne et al. 2020, Vizentin-Bugoni et al. 2014). For example, in bird–plant interactions, interactions are influenced by the match between fruit diameter and bill width, which determines whether a fruit can be used by a frugivore. However, so far, analyses on the influence of functional traits on species interactions have mostly been restricted to matching of individual trait pairs (Burns 2013; Dehling et al. 2014b) or the matching of trait combinations (Dehling et al. 2016), and even the latter are limited to relating the morphology of a species (its own trait combination) to a single-point representation of its niche, for instance, a species’ niche position (determined as the mean value of the trait combinations of their different interaction partners). Our proposed approach of simultaneously considering multiple traits in a single joint model facilitates the consideration of the entire range of trait combinations that a consumer species uses and which, in their entirety, describe the species’ Eltonian niche.

Modelling the probability of consumer–resource interactions via the joint modelling of Eltonian niches can help improve the modelling of responses for species with few observed interaction partners, for instance, for those species that are naturally rare or difficult to observe in the field. As we showed in our example, due to the strong relationship between the birds’ functional traits and their Eltonian niches, it was even possible to accurately predict interactions for species for which there are *no* empirical observations available— based entirely on their similarity in functional-trait values with the other bird species in the network. This demonstrates the power and main advantage of the joint modelling approach: empirical interaction networks are notoriously sparsely sampled (Jordano 2016), and—as revealed by studies comparing networks compiled from direct observations with networks sampled indirectly using molecular techniques (Pornon et al. 2017)—a large proportion of interactions in ecological networks are readily missed. The ability to predict unobserved interactions will lead to more complete data on species interactions which might lead to a better understanding of the drivers underlying interactions in ecological communities. Predicting new interactions from proxies is especially valuable when attempting to model new interactions between species that have never interacted before, for example in communities with altered composition due to climate-change induced range shifts or the introduction of exotic species. This facilitates assessing the potential impact of environmental changes and invasive species on ecological communities.

Our example in which we applied the model to bird–plant seed-dispersal interactions was based on previous studies that showed that interactions between plants and frugivorous birds are influenced by functional traits (Dehling et al. 2014b, 2016; Eklöf et al. 2013). It will be interesting to test our approach to modelling Eltonian niches in the context of other interactions. We expect that our model might work best with other types of interactions that are influenced by the traits of the interacting partners, such as plant–pollinator interactions (Peralta et al. 2020b; Sonne et al. 2020; Vizentin-Bugoni et al. 2014) or predator–prey interactions that are determined by size relationships (Gravel et al. 2014). The approach might be less powerful in generalistic systems or taxa (e.g. species in extreme or disturbed environments), where trait matching tends to be weaker (Peralta et al. 2020a).

However, it is also possible that relationships between species’ functional traits and their Eltonian niches have been overlooked in such systems. The increased availability of a wider set of functional traits might eventually lead to the identification of trait-matching relationships especially in those types of networks that are currently understudied. Our Eltonian niche models can serve as a tool to identify the functional traits that influence species interactions across different types of networks.

### Further improvements of Eltonian niche modelling

Even though the explanatory and predictive power of our models were high, the usability of our niche models could still be improved by further disentangling ecologically feasible interactions (fundamental Eltonian niches) from locally realized interactions (realized Eltonian niches). As species interactions are usually studied on the level of species communities, this could lead to the assumption that all interactions between suitable co-occurring partners are realized; however, restrictions to species interactions might occur on smaller scales and could be difficult to sample in the field. For instance, even though two suitable interaction partners are present in a local community, their interaction might ultimately depend on whether the home ranges (or growing sites) overlap or whether both species occur in sufficient numbers to encounter each other. Indeed, local abundances can easily influence whether a consumer interacts with a suitable resource (Martins et al. 2024). For instance, at a certain site, a species might not use a potential resource as long as its preferred resource (e.g. the one to which it is best adapted) is abundantly available (hence, using a narrower range of resources), whereas it might shift to less preferred resources at sites where its preferred resource is rare.

In the absence of such detailed information, our approach of using empirical data and known trait relationships underlying species interactions to restrict absences to the extremely unlikely or impossible interactions (forbidden links), while treating possible, but unobserved, interactions as NA therefore served as a cautious and conservative compromise. Nevertheless, the inclusion of local species abundances into the modelling framework might facilitate additional insights into observed differences in interaction probabilities between suitable interaction partners and into species’ preferences for certain resources within their Eltonian niche (Bender et al. 2018, Coux et al. 2020, Martins et al. 2024). Similarly, in our example of plant–frugivore interactions we could not consider intraspecific differences in resource use (simply because sampling the resource use of individual birds is virtually impossible in empirical plant–frugivore interactions). However, the inclusion of more-highly resolved networks on the level of individuals (Guimarães 2020; Hervías-Parejo et al. 2020; Rumeu et al. 2018; Wells and O’Hara 2013), potentially aided by the use of automatic detection (Weinstein 2015) or indirect sampling of interactions via molecular techniques (reviewed in Pringle & Hutchinson 2020), and combined with individual-level measurements of functional traits for consumers and resources, might lead to further insights into intraspecific variation in Eltonian niches and the mechanisms underlying the restrictions of locally realized interactions.

In addition to using methods derived for the modelling of species’ Grinnellian niches (JSDMs) to study species’ Eltonian niches, as demonstrated in this article, we recognize that there are also likely benefits to be obtained from exploring research questions related to Grinnellian and Eltonian niches together (rather than either or). For instance, data on species’ Eltonian niches (the range of resources used by each species) could be integrated into Grinnellian-niche models to improve the modelling of species ranges by explicitly capturing dependences on the distribution of suitable resources. The consideration of species’ responses to both environmental conditions and available resources will likely lead to an improved understanding of species’ potential range shifts and the composition of future species assemblages in response to environmental changes (Dormann et al. 2020).

## Acknowledgements

DMD acknowledges support by ARC SRIEAS Grant SR200100005 Securing Antarctica’s Environmental Future. HRL acknowledges support by the Marsden Fund managed by the Royal Society Te Apārangi (grant MFP-UOC2102) and the Bioprotection Aotearoa Centre of Research Excellence.

## Box 1: Disentangling observed from unobserved interactions

The modelling of niches using a joint hierarchical model, such as JSDM, requires robust information about presence *as well as* absence data. For the modelling of Grinnellian niches, such data are usually readily available because species occurrences are comparatively well-known. In addition, Grinnellian niches are commonly modelled based on the environmental conditions found within a species’ range (either continuous ranges or a set of point localities) and based on the assumption that these conditions are continuous in space (Renner et al. 2015). This has the advantage that even for the rarest species it is usually possible to determine a range of environmental conditions (e.g., at least the seasonal variation in the type locality) that can be used to model its niche.

However, using the same approach and assumptions for interaction data is usually not possible. This is because species interactions are much more difficult to record than species occurrences, and the sampling of interaction networks therefore tends to be notoriously sparse and opportunistic (Jordano 2016). Moreover, the presence of an interaction is much more reliably sampled than the absence and, consequently, there is high uncertainty with regard to the proportion of unobserved interactions per species. In the same sense as geographic ranges, network data are more akin to sparsely-sampled point occurrences.

The requirements for the modelling of Eltonian niches are therefore slightly different from those for Grinnellian niches. Assuming that a consumer can interact with all resources that show similar trait combinations as its observed resources—analogous to drawing ranges that connect known point occurrences—would likely work relatively well for abundant species because their resources are well represented in a sample; in contrast, it would probably underestimate the size of niches for species that are rare or difficult to observe, and for which therefore only few observations are available that are less likely to be a good representation of their niches. Treating all missing interactions as absences (structural zeros), however, likewise represents an unrealistic assumption because rather than a true absence (e.g. either an impossible interaction or the active avoidance of a resource), missing interactions include many interactions that were simply not observed (Blasco-Moreno et al. 2019).

To separate unobserved interactions from true absences for each species, we calculated the ratio between the plant’s fruit diameter and the bird’s bill width—a well-known trait-matching relationship in bird–plant interactions (Dehling et al. 2014b)—across all observed interactions in the empirical network (398 species pairs, coded as “1” in the matrix), and then used the inner 95th percentile of this ratio (range 0.2–1.5) as limits for unobserved, but possible interactions, which is a conservative estimate. We then used these limits to separate the 2,774 unobserved interactions into those considered possible (2,370 species pairs, coded as “NA” in the matrix) and those considered unlikely or impossible (“forbidden links”; 404 species pairs, coded as “0” in the matrix), following Lai et al. (2021). Another analysis treating all missing interactions as true absences (i.e., without NAs) yielded qualitatively similar results (Appendix 1, Table S1).

## Appendix 1

**Table S1.** Explanatory and predictive power, explained variance, and phylogenetic signal in Eltonian niches of four joint hierarchical models describing the interactions between 61 frugivorous birds and 52 fleshy-fruited plants. Eltonian niches of bird species are described by the functional traits of the plants with which the birds interact. The joint hierarchical models relate the birds’ Eltonian niches to different combinations of co-variates (no co-variates, bird traits, bird phylogeny, bird traits + bird phylogeny). Percent variance explained describes the mean variance in bird–plant interactions explained by plant traits and the mean variance in Eltonian niches explained by bird traits. The phylogenetic signal in the birds Eltonian niches is given as the 95% interval and median value.

| Model | Co-variables | WAIC | Explanatory power |  | Predictive power |  | Percent variance explained |  | Phylogenetic signal<br>bird traits |
| --- | --- | --- | --- | --- | --- | --- | --- | --- | --- |
|  |  |  | AUC | Tjur R<sup>2</sup> | AUC | Tjur R<sup>2</sup> | bird-plant interactions<br>by plant traits | Eltonian niches<br>by bird traits |  |
| 1 | no co-variables | 16.64 | 0.9 | 0.25 | 0.56 | 0.02 | 81.9 | - | - |
| 2 | bird traits | 16.63 | 0.9 | 0.26 | 0.60 | 0.03 | 83.4 | 42.2 | - |
| 3 | bird phylogeny | 16.59 | 0.9 | 0.24 | 0.55 | <0.01 | 77.3 | - | 0.66–1 (0.91) |
| 4 | bird traits + bird phylogeny | 16.53 | 0.9 | 0.25 | 0.54 | <0.01 | 79.8 | 45.3 | 0.63–1 (0.91) |

**Fig. S1.**
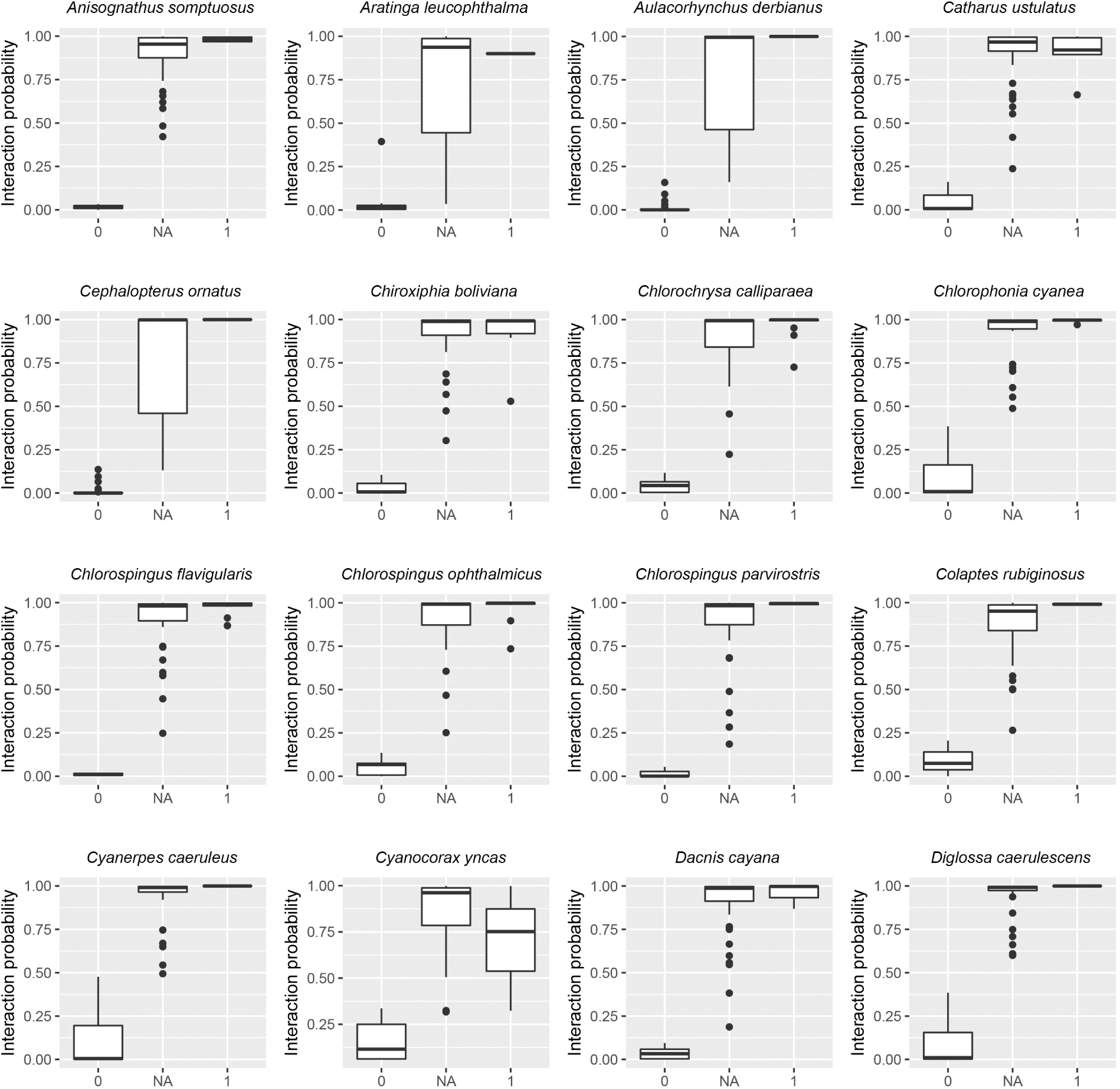
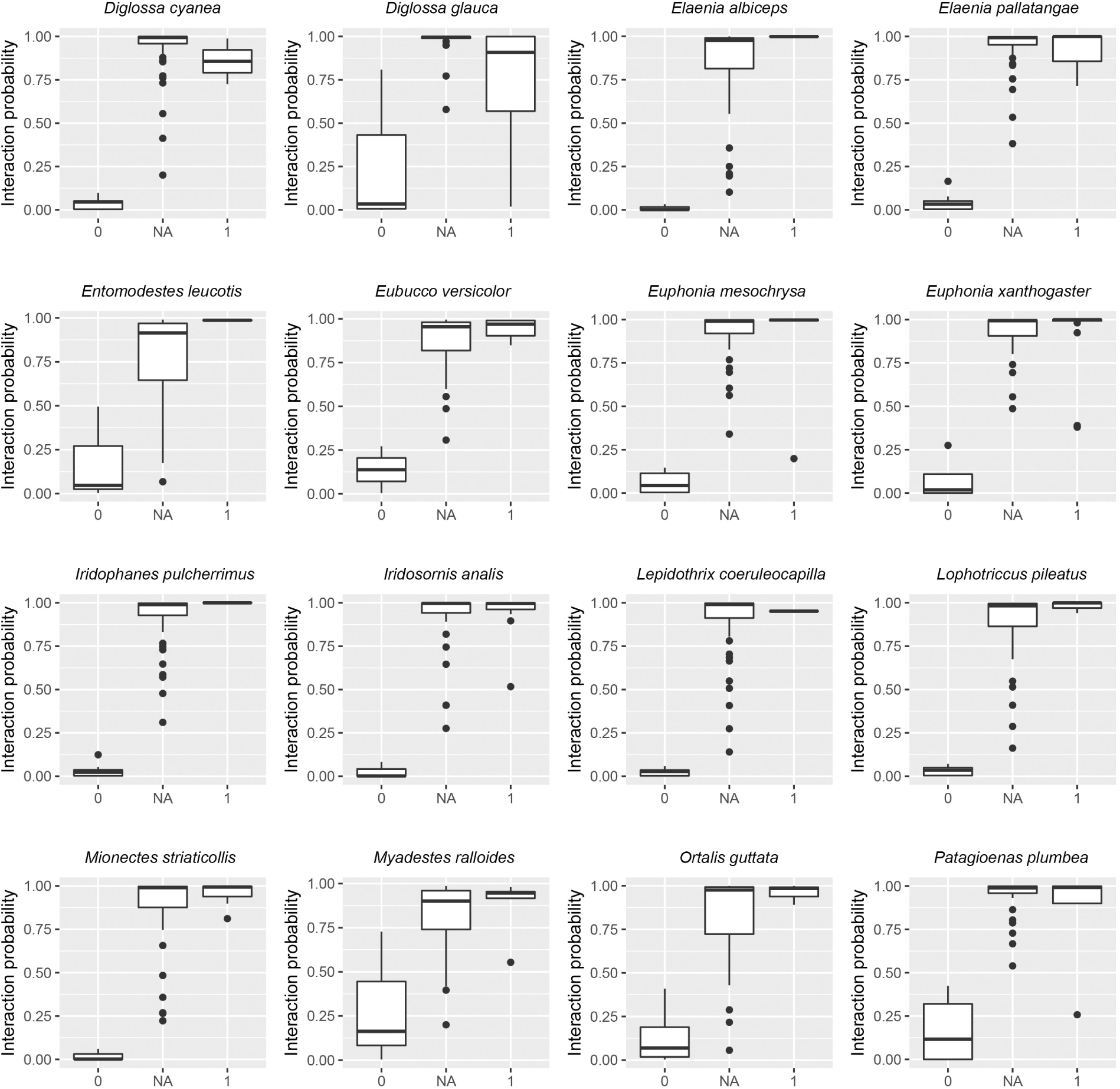
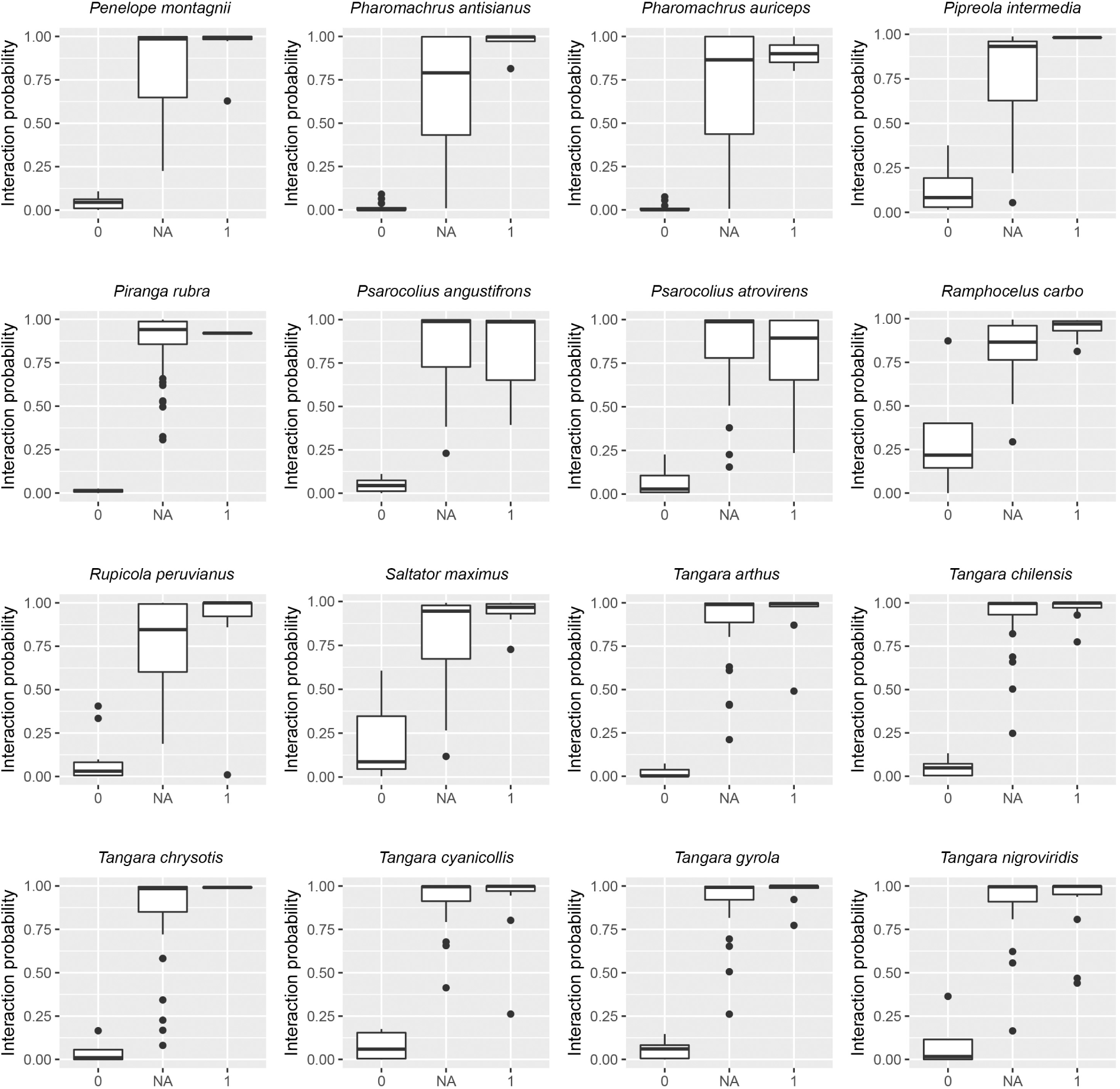
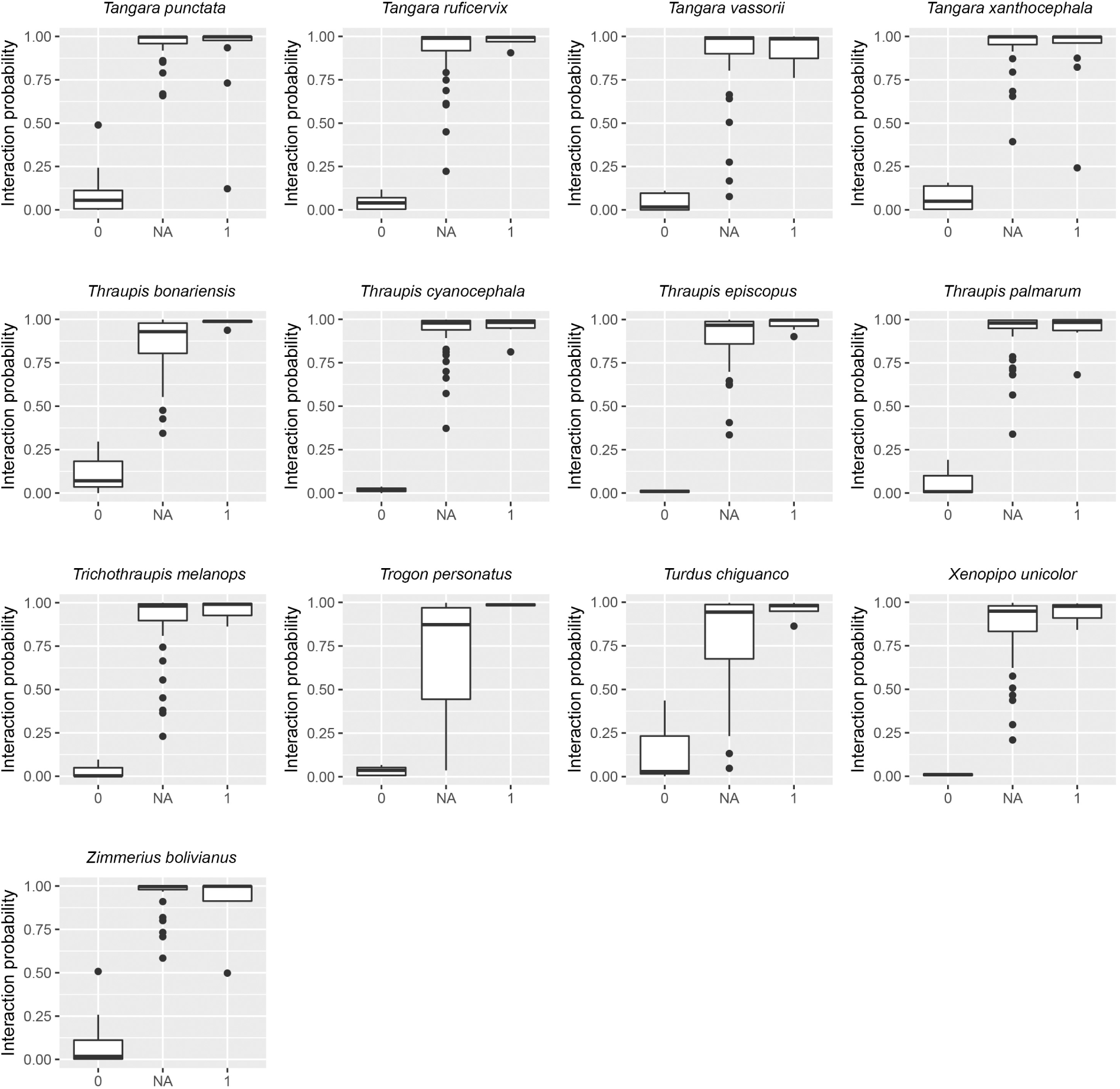
Interaction probability for 61 bird species modelled with a joint hierarchical niche model (model 2) based on an empirical network of bird–plant interactions from the tropical Andes. Interactions in the network were marked as observed (1), unobserved but likely (NA), and unobserved and unlikely (0); see Methods for details.

**Fig. S2.**
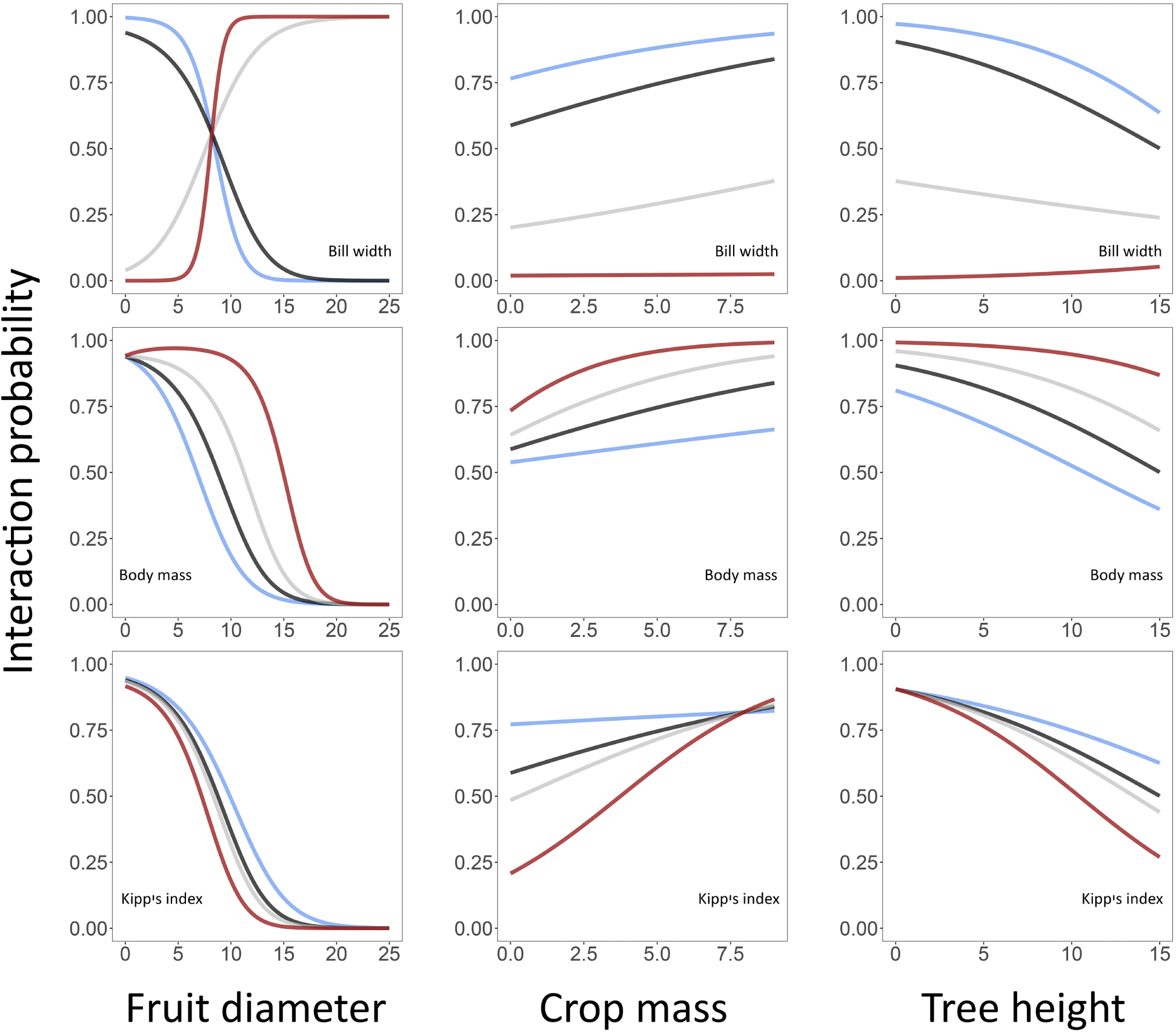
Interaction probabilities between frugivorous birds and fleshy-fruited plants determined by bird and plant traits. Bird traits (rows) include bill width, body mass, and Kipp’s index; plant traits (columns) include fruit diameter, crop mass, and tree height. Posterior-mean interaction probabilities predicted by model 4 (*y*-axis) are shown along a gradient of a focal plant trait (*x*-axis) for four hypothetical bird species with different values of a focal bird trait (blue, 0.025 percentile; black, 0.5 percentile; grey, midpoint between the 0.025 and the 0.975 percentiles; red, 0.975 percentile). For all predictions, non-focal plant traits and non-focal bird traits were set to the respective community median.

**Fig. S3.**
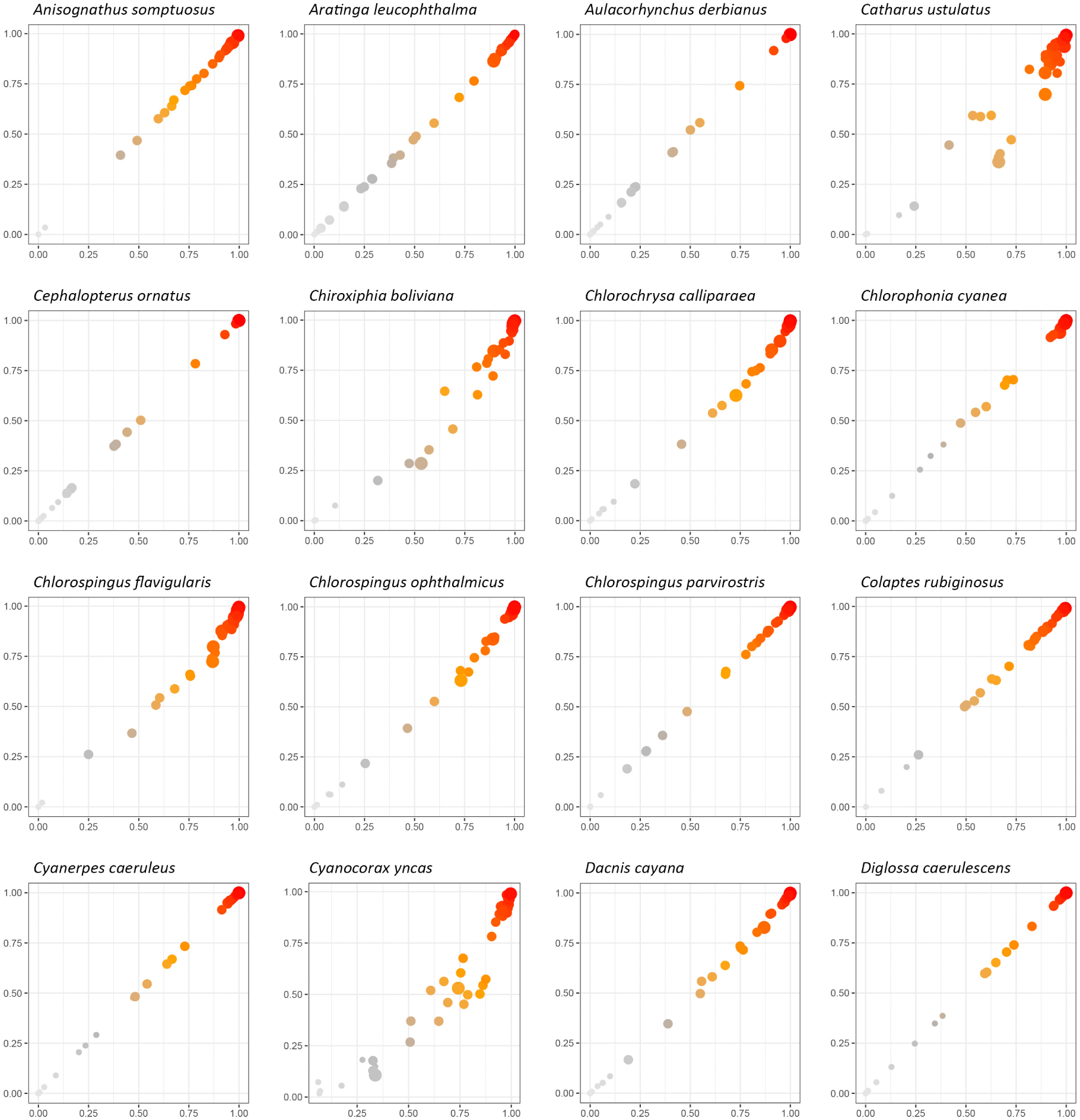
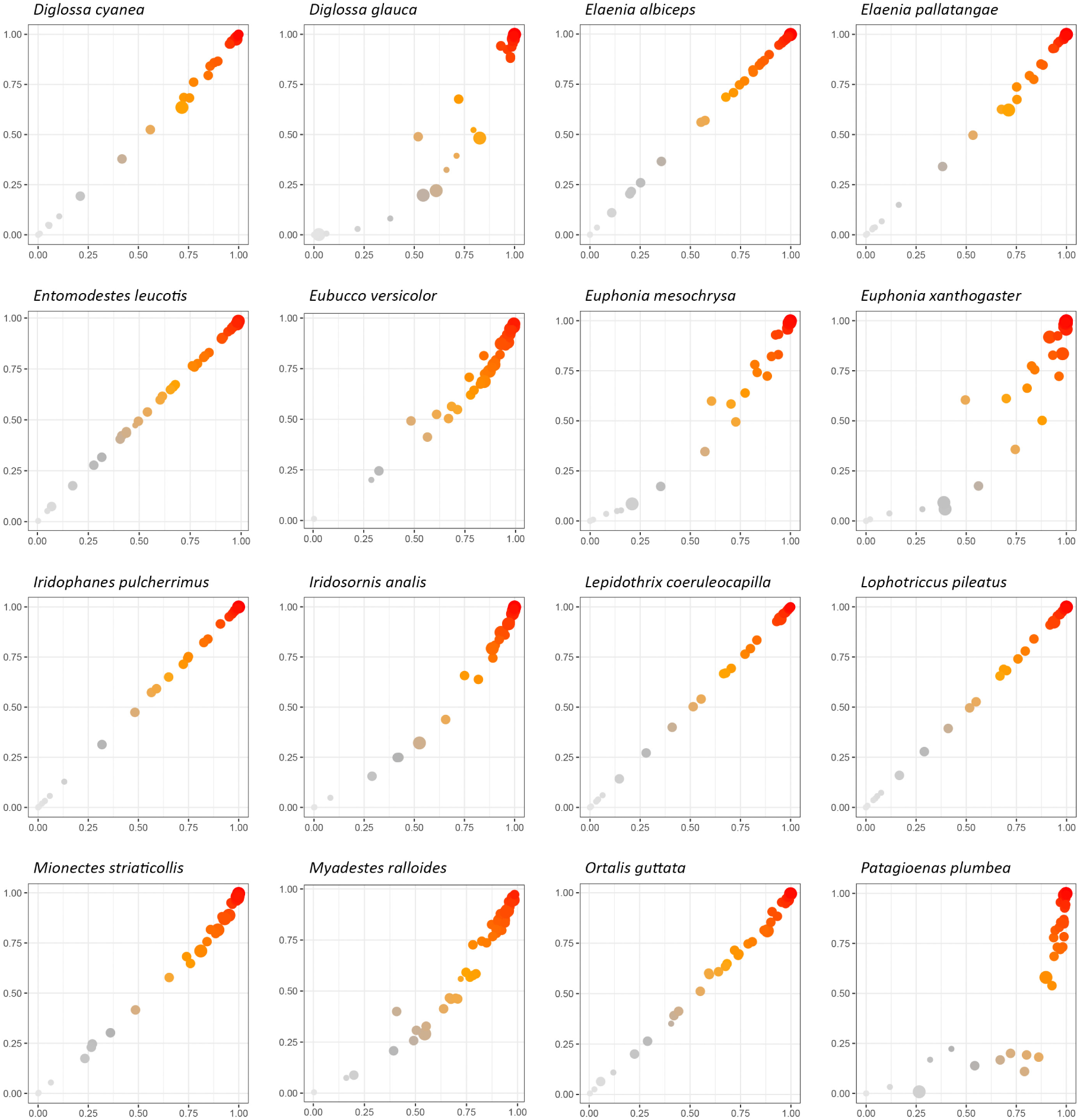
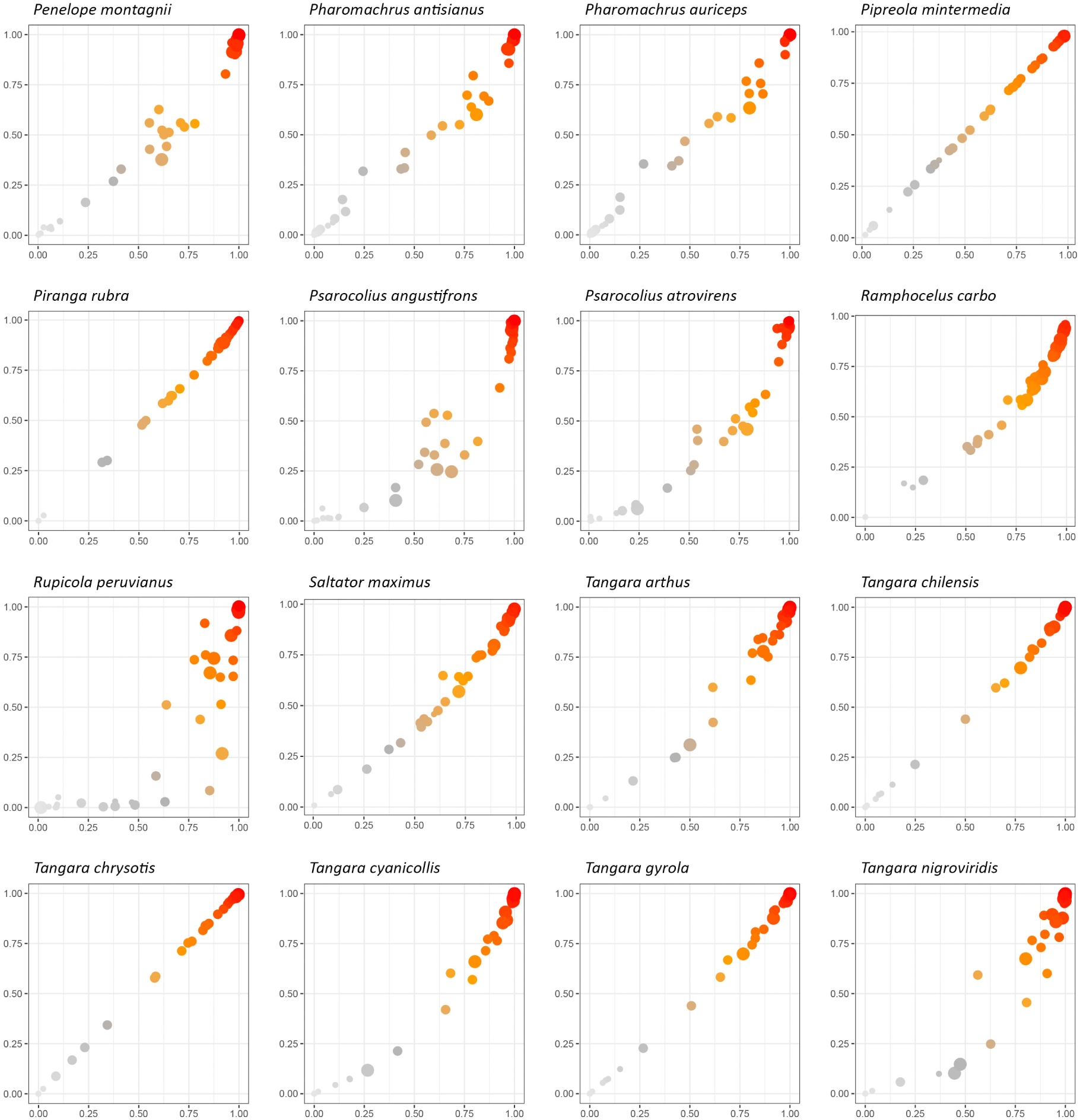
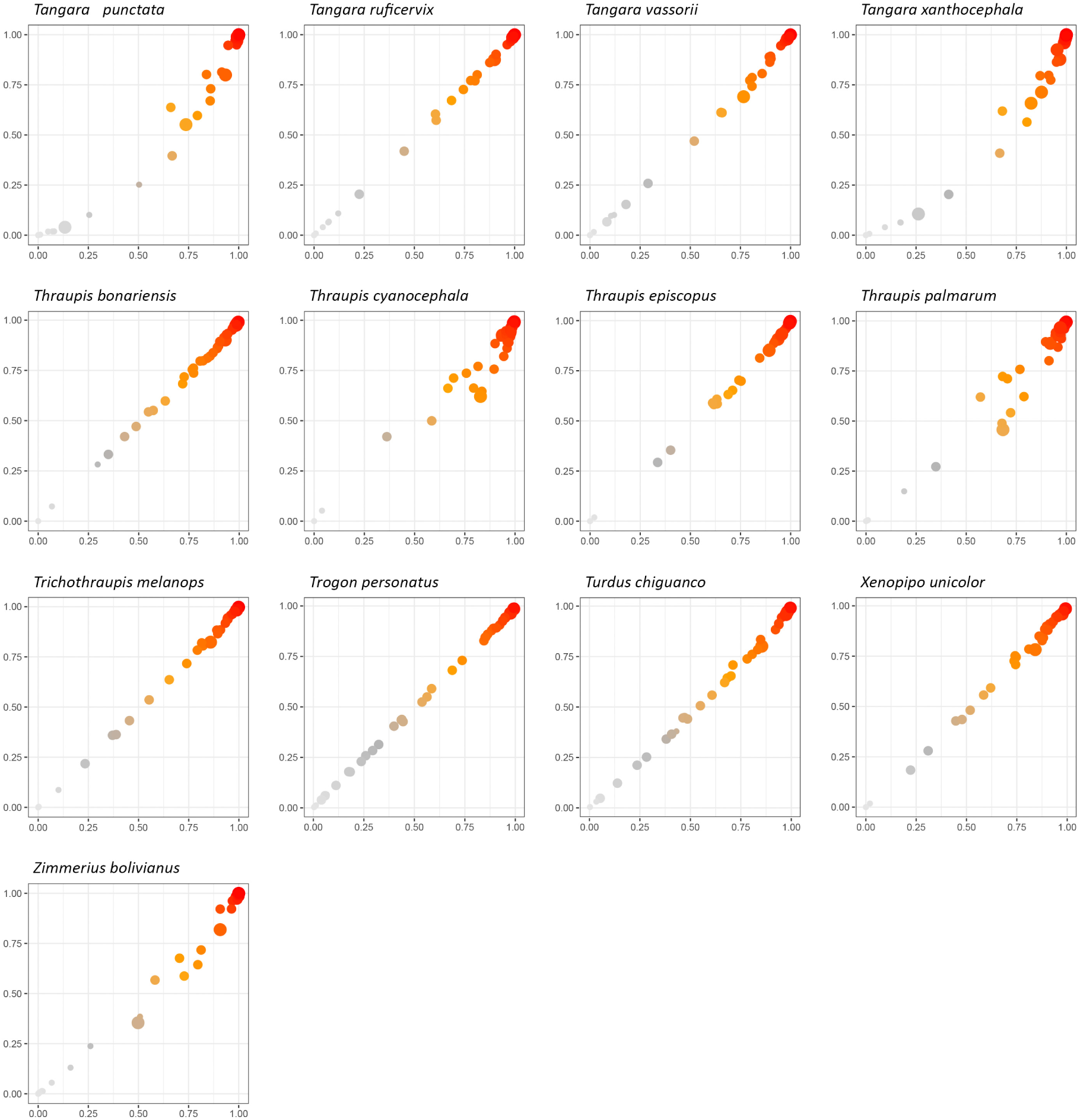
Correlations between the interaction probabilities modelled from observed interactions and those predicted based on birds’ functional traits for each bird species in the network.

## Appendix 2

In our maximal model 4, we modelled the presence–absence of interaction, *Y_ij_*, between plant (resource) species *i* and bird (consumer) species *j* in a joint hierarchical model as:

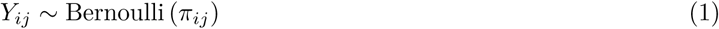

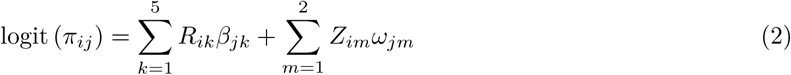

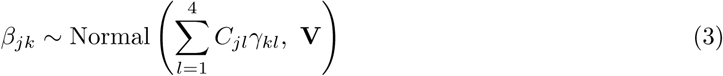

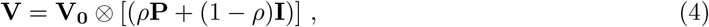

where *π_ij_* is the probability of interaction between plant species *i* and bird species *j*. The intercept *β_j_*_1_ (that corresponds to the intercept column *R_i_*_1_ = 1 ∀ *i*) accounts for interspecific variation in average interaction probability across bird species. The consumer-specific slopes *β_j,k>_*_1_, which can be interpreted as the bird species’ Eltonian niche, estimate bird species’ responses to the three plant resource traits *R_i,k>_*_1_. We also included two sets of plant- and bird-specific latent variables, *Z_i_* and *ω_j_* respectively, to account for co-variation in bird species’ resource use. Next, we allowed the three bird consumer traits *C_jl_* (plus an intercept column *C_j_*_1_ = 1 ∀ *j*) to moderate their Eltonian niche *β_jk_* via the trait–trait interaction term, *γ_kl_*. Lastly, the term **V** captures any unexplained variation in Eltonian niche not explained by bird traits, and is further partitioned into an overall variation unrelated to bird phylogeny, **V_0_**, and another component that is potentially influenced by bird phylogeny **P** by a proportional factor 0 *< ρ <* 1; **I** is an identity matrix necessary for the estimation of phylogenetic signal in Eltonian niche.

In model 3, we omitted bird traits by removing the *β_j,k>_*_1_ and *R_i,k>_*_1_ terms. In model 2, we omitted bird phylogeny by setting **V** = **V_0_** (or equivalently *ρ* = 0). In model 1, we performed both of these omissions.

